# Site of breast cancer metastasis is independent of single nutrient levels

**DOI:** 10.1101/2024.10.24.616714

**Authors:** Keene L. Abbott, Sonu Subudhi, Raphael Ferreira, Yetiş Gültekin, Sophie C. Steinbuch, Muhammad Bin Munim, Sophie E. Honeder, Ashwin S. Kumar, Diya L. Ramesh, Michelle Wu, Jacob A. Hansen, Sharanya Sivanand, Lisa M. Riedmayr, Mark Duquette, Ahmed Ali, Nicole Henning, Anna Shevzov-Zebrun, Florian Gourgue, Anna M. Barbeau, Millenia Waite, Tenzin Kunchok, Gino B. Ferraro, Brian T. Do, Virginia Spanoudaki, Francisco J. Sánchez-Rivera, Xin Jin, George M. Church, Rakesh K. Jain, Matthew G. Vander Heiden

## Abstract

Cancer metastasis is a major contributor to patient morbidity and mortality^1^, yet the factors that determine the organs where cancers can metastasize are incompletely understood. In this study, we quantify the absolute levels of over 100 nutrients available across multiple tissues in mice and investigate how this relates to the ability of breast cancer cells to grow in different organs. We engineered breast cancer cells with broad metastatic potential to be auxotrophic for specific nutrients and assessed their ability to colonize different organs. We then asked how tumor growth in different tissues relates to nutrient availability and tumor biosynthetic activity. We find that single nutrients alone do not define the sites where breast cancer cells can grow as metastases. Additionally, we identify purine synthesis as a requirement for tumor growth and metastasis across many tissues and find that this phenotype is independent of tissue nucleotide availability or tumor de novo nucleotide synthesis activity. These data suggest that a complex interplay of multiple nutrients within the microenvironment dictates potential sites of metastatic cancer growth, and highlights the interdependence between extrinsic environmental factors and intrinsic cellular properties in influencing where breast cancer cells can grow as metastases.

## Introduction

Understanding the factors that govern tumor growth in metastatic sites is important for developing more effective therapies for advanced cancer. Many factors in the tumor microenvironment contribute to where tumors can grow as metastases, with nutrient availability being one important component^2–9^. Variations in nutrient availability across different tissues can limit the sites where cancers can grow as metastases^3,4,10,11^, arguing specific metabolic adaptations are required for cancer cells to colonize a metastatic tissue site.

Low availability of select nutrients in some tissues can impose metabolic constraints that influence tumor growth and metastatic potential^10,11^. For instance, limited lipid availability in the central nervous system microenvironment results in a site-specific dependency on fatty acid synthesis or fatty acid desaturation for the metastasis of breast tumors^12,13^ or leukemias^14^ to this tissue. Similarly, reduced serine levels in the brain have been shown to cause breast cancer brain metastases to become vulnerable to inhibition of the serine synthesis enzyme PHGDH^15^. In the lungs, which are rich in lipids, targeting palmitate processing via CPT1a can inhibit breast cancer-derived lung metastasis^16^. Both pyruvate and asparagine availability have also been found to influence breast cancer metastasis to the lungs^17,18^. These studies raise the question of whether reduced availability of specific nutrients in certain tissues broadly predicts where metastatic cancer cells are likely to colonize and grow.

Various methods have been developed to experimentally model metastasis in mice, including development of a first-generation metastasis map using an in vivo barcoding strategy^12^. This approach evaluated the metastatic potential of 493 human cancer cell lines across five organs via intracardiac injection and was used to uncover the role of altered lipid metabolism in breast cancers that metastasize to the brain. Inspired by this method, we sought to explore the relationship between tissue nutrient availability and metastatic potential in triple-negative breast cancer (TNBC). To achieve this, we quantified the absolute levels of 112 metabolites present across seven mouse tissues and constructed a series of nutrient auxotrophs to assess their metastatic potential to multiple tissues following intracardiac injections. Our analysis revealed that while levels of some metabolites correlate with metastatic potential, the levels of individual nutrients in isolation are insufficient to determine metastatic preference or the ability of specific nutrient auxotrophs to grow in a tissue site. Instead, our findings suggest that metastatic preference is driven by a combination of multiple nutrient levels and cell-intrinsic metabolic factors.

## Results

### Nutrient availability differs across tissues

To broadly assess nutrient availability across multiple tissue sites, including where TNBC cells can grow as primary tumors or metastases, we isolated plasma and interstitial fluid (IF) from the mammary fat pad (MFP), liver, lung, kidney and pancreas of NOD-SCID-gamma (NSG) mice. We also collected cerebrospinal fluid (CSF) as a surrogate for the brain extracellular environment because acquiring sufficient IF for analysis of multiple metabolites from normal brain tissue was not feasible. We then conducted quantitative mass spectrometry to determine the absolute levels of 112 metabolites in the IF from these different tissue sites (Fig. 1a-c, Extended Data Fig. 1a-j) and confirmed high correlations between metabolite concentrations measured in IF isolated from the same sites from two independent NSG mouse cohorts (Extended Data Fig. 1k). Principal component analysis (PCA) and hierarchical clustering revealed that metabolites measured in tissue IF samples cluster distinctly from those measured in plasma and CSF (Fig. 1b-c). When considering the differences in metabolite concentration between tissue IF and plasma, we found that while some metabolites were depleted in tissue IF relative to plasma, numerous metabolites were present at higher concentration in IF than plasma (Fig. 1d). In contrast, CSF showed lower levels of many metabolites compared to plasma, a pattern likely attributed to the selective permeability of the blood-brain barrier and that is consistent with previous studies examining both humans and mice^19,20^. Notably, we found that nucleotide-related metabolites, but not amino acids, were greater contributors to the PCA components separating the fluid samples (Fig. 1e). This finding suggests that levels of nucleotides and nucleotide precursors/salvage products are an important contributor to the differences in nutrient availability across tissue environments.

**Fig. 1:**
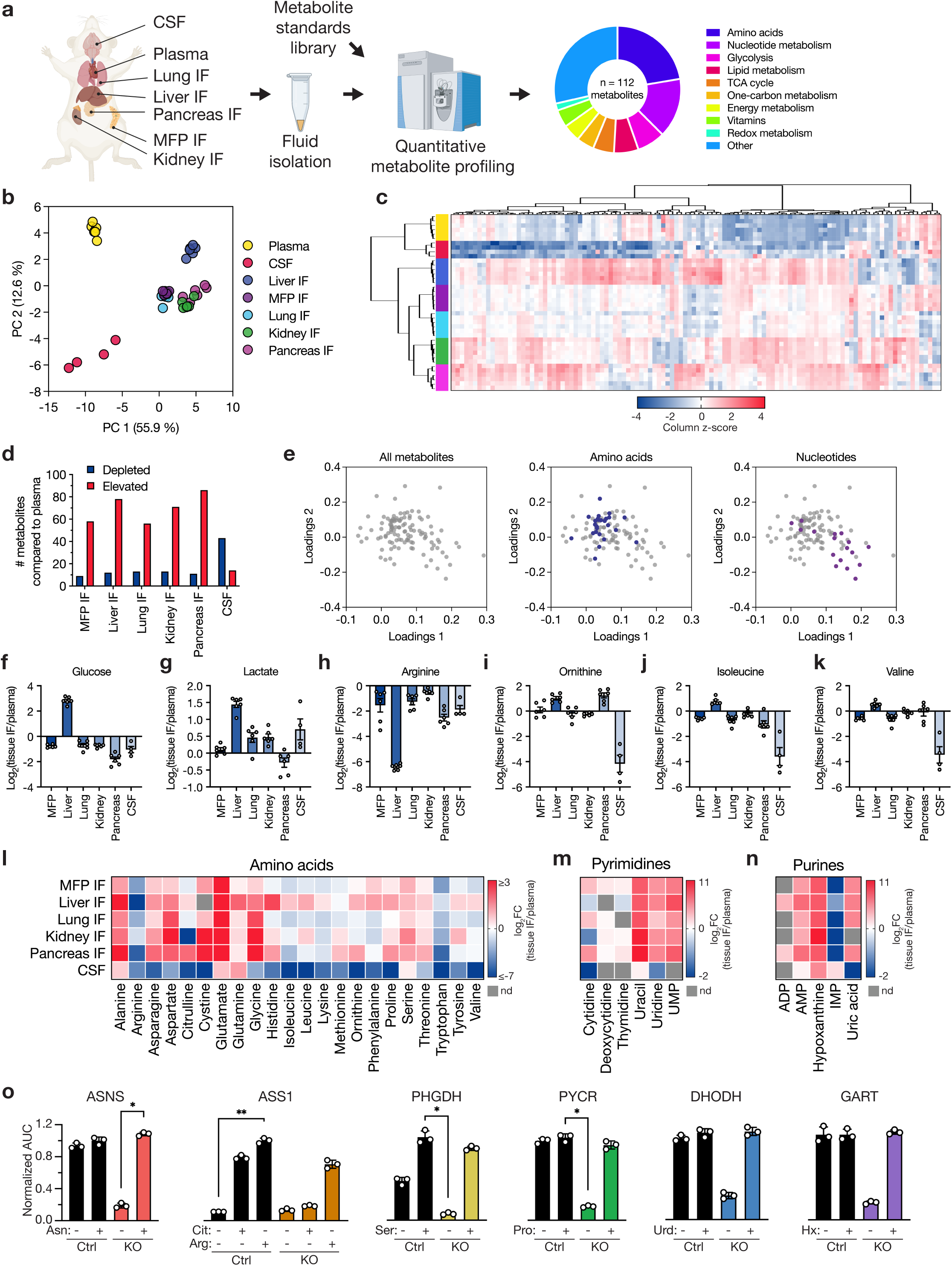
Nutrient levels in tissue interstitial fluid, plasma, and CSF from mice. **a**, Schematic depicting the isolation and quantification of metabolites from plasma, cerebrospinal fluid (CSF), and tissue interstitial fluids (IF) from female NOD-SCID-gamma (NSG) mice. Metabolites in these fluid samples were measured using LC/MS, with quantification performed alongside a dilution series of chemical standards. In total, 112 metabolites were identified and quantified across the different samples. MFP: mammary fat pad. **b-c**, Principal component analysis (PCA) (b) or hierarchical clustering (c) of metabolites measured in tissue IF samples, plasma, and CSF. Data represent n = 6 (plasma, kidney IF, liver IF, lung IF, MFP IF, pancreas IF) or n = 4 (CSF) biological replicates. Data presented within each column of the heatmap were z score normalized. **d**, Bar plot showing the number of metabolites with significantly lower (depleted) or higher (elevated) concentrations in each tissue IF or CSF relative to plasma. A fold change of 2 and a raw p-value of 0.05 assuming unequal variance were used to select significantly altered metabolites based on a t-test analysis. **e**, Loadings plot presenting the contribution of individual metabolites to the PCA components in (b). Metabolites are colored according to their assignment to amino acid or nucleotide metabolism pathways as indicated. **f-k**, log2 fold change in the indicated metabolite concentrations measured in each tissue IF or CSF relative to plasma. Data are presented as mean ± SEM and represent n = 6 (kidney IF, liver IF, lung IF, MFP IF, pancreas IF) or n = 4 (CSF) biological replicates. **l-n**, Heatmaps depicting the average log2 fold change in the indicated metabolite concentrations measured in each tissue IF or CSF relative to plasma. Scale bars provided for each group indicate the ranges of values shown. **o**, Area under the curve (AUC) values derived from proliferation curves for MDA-MB-231 control or knockout (KO) cells cultured with or without the relevant rescue metabolites presented in Extended Data Fig. 5. AUC values were normalized to the control cell line treated with the relevant rescue metabolite. Data are mean ± SD and represent n = 3 biological replicates and statistical analysis was performed using a Kruskal-Wallis test with Dunn’s multiple comparisons test (*p < 0.05, **p < 0.01). Arg: arginine; Cit: citrulline; Ser: serine; Pro: proline; Urd: uridine; Hx: hypoxanthine. PYCR represents PYCR1, 2, and 3 triple KO.

Many of the differences in metabolite concentrations across tissues align with known metabolic characteristics of each tissue. Most tissue IF displayed lower glucose and higher lactate levels than plasma (Fig. 1f-g), consistent with active glucose metabolism^21^. The liver stood out with higher levels of both glucose and lactate in IF compared to plasma, consistent with its glycogenolytic and gluconeogenic capabilities, as well as its role in recycling lactate back to glucose^22^. In liver IF, we also observed lower arginine and higher ornithine levels than plasma (Fig. 1h-i), likely reflecting active arginase activity in this tissue^23^. Conversely, citrulline was depleted in kidney IF (Extended Data Fig. 2a), aligning with human data^24^ and perhaps reflective of an active urea cycle in this tissue. Liver IF contained relatively higher concentrations of branched-chain amino acids (BCAAs: isoleucine, leucine, valine) than other tissues (Fig. 1j-k, Extended Data Fig. 2b), which may reflect known low BCAA aminotransferase activity in this organ relative to other tissues^25^.

Consistent with previous measurements^19,20^, many amino acids were lower in CSF compared to circulation, with notable variation in amino acid levels measured in other tissues (Fig. 1l, Extended Data Fig. 1a). Unlike previous reports^19,20^, serine was not depleted in the CSF relative to plasma, though it was lower compared to other tissue IF. This lack of serine depletion in CSF was consistent across both NSG and C57BL/6J mouse strains (Extended Data Fig. 2c), and may be influenced by other factors such as circadian rhythm^26^ or diet^27^. Additionally, while many nucleotide species were elevated in tissue IF and low in CSF compared to plasma (Fig. 1m-n, Extended Data Fig. 1d, 2d), the purine hypoxanthine was consistently high across all tissue IF as well as in CSF. Collectively, these findings denote many variances in nutrient availability across different tissues, which could constrain the ability of cancer cells to metastasize and grow in particular organ sites.

### Generation of breast cancer cells auxotrophic for specific amino acids or nucleotides to study the relationship between nutrient auxotrophy and metastatic site

Cancer cells exposed to nutrient-limited microenvironments often adapt by upregulating metabolic pathways to synthesize the deficient metabolite, ensuring their survival and proliferation^10^. To eliminate this adaptive response as a confounding factor and isolate how nutrient availability constrains metastatic growth, we engineered widely metastatic TNBC cell lines to be auxotrophic for specific metabolites that vary across tissues. We used the human lines MDA-MB-231 and HCC1806^12^, as well as the murine-derived line EO771^28^, and employed CRISPR/Cas9 to knock out genes essential for the synthesis of specific metabolites with known nutrient rescues (Extended Data Fig. 3a). Specifically, we targeted *ASNS* (asparagine^29,30^), *ASS1* (arginine^31,32^), *PHGDH* (serine^15,33^), *PYCR1,2,3* (proline^34^), *DHODH* (pyrimidines, e.g. uridine^35,36^), and *GART* (purines, e.g. inosine or hypoxanthine^37^). We confirmed variable expression of each of these proteins in parental MDA-MB-231, HCC1806, and EO771 cells by western blot (Extended Data Fig. 3b-c); notably, MDA-MB-231 cells expressed lower levels of ASS1 and PHGDH, consistent with prior observations^33,38^. We also confirmed loss of relevant enzyme expression in each knockout line by western blot (Extended Data Fig. 4), and further confirmed that each knockout line exhibited impaired proliferation only in the absence of the relevant nutrient that corresponded to the intended auxotrophy (Fig. 1o, Extended Data Fig. 5). Importantly, supplementation of the specific metabolite rescued proliferation of each auxotroph cell line to comparable levels as the parental line. In order to generate proline auxotrophs, deletion of the mitochondrial (*PYCR1,2*) and the cytoplasmic (*PYCR3*) genes encoding pyrroline-5-carboxylate reductase was required, as the presence of any of these genes has been reported to enable proline synthesis^34^ and support proliferation following proline withdrawal (Extended Data Fig. 4d, 5d). Since arginine starvation in parental cells arrests proliferation via mTOR inactivation^39,40^, addition of citrulline was required for control cells to proliferate without arginine (Fig. 1o, Extended Data Fig. 5b); however, *ASS1*-null cells could not grow without arginine even if citrulline was added, validating that *ASS1* knockout cells lost the ability to generate arginine from citrulline. For GART, despite the presence of some species detected at 110 kDa^41^ in MDA-MB-231 and HCC1806 using an anti-GART antibody (Extended Data Fig. 4f), depletion of the 50 kDa monomeric form was sufficient to result in hypoxanthine auxotrophy in all three cell lines (Extended Data Fig. 5f). Collectively, these data confirm that the genetic modifications render cells dependent on external supplementation of the expected metabolites for proliferation and survival in standard culture conditions.

In constructing the nutrient auxotrophs, we identified three distinct categories of metabolite requirements. First, removing asparagine or proline from the culture medium reduced cell proliferation only when the relevant synthesis genes were knocked out (Extended Data Fig. 5a, d). Second, withdrawal of arginine in all cell lines, or serine in MDA-MB-231 cells, reduced proliferation even when the synthesis genes were intact (Extended Data Fig. 5b, c). Third, nucleotides, which are typically absent from standard culture media, could be supplemented to rescue cell dependency on *DHODH* or *GART* (Extended Data Fig. 5e, f). These findings indicate that for some nutrients, cellular synthesis capacity is important for maximal proliferation regardless of environmental availability. Nevertheless, this panel of metabolite auxotrophs with varying metabolite synthesis requirements enable testing of how nutrient levels influence the ability of cells to colonize different tissues.

### Availability of single nutrients does not predict the tissues where breast cancer cells can grow as metastases

Prior to using the different auxotrophic cells to study how tissue nutrient availability influences tumor growth in different metastatic sites, we first validated the metastatic potential of control MDA-MB-231, HCC1806, and EO771 cells to grow as tumors in different sites. We engineered each line to express firefly luciferase (Fluc), then injected the line into the left ventricle of NSG mice (MDA-MB-231, HCC1806 cells) or C57BL/6J mice (EO771 cells) and monitored tumor growth over time via bioluminescence imaging (Extended Data Fig. 6a). At endpoint, we harvested brain, liver, lungs, bone, kidney/adrenal gland, and ovaries, and quantified bioluminescence in each tissue to determine tumor burden (Extended Data Fig. 6b-c). This verified the ability of these cells to metastasize to multiple tissues, aligning with the previously reported metastatic potential of MDA-MB-231 and HCC1806 cells^12^.

Next, we performed intracardiac injections of each Fluc-tagged auxotroph or control cell line and assessed tissue-specific bioluminescence of the auxotrophs relative to the control lines three weeks post-injection for EO771 cells and four weeks post-injection for MDA-MB-231 or HCC1806 cells (Fig. 2a). From these experiments, we found that nucleic acid auxotrophs show an overall consistent impairment in their ability to grow in multiple tissues (Fig. 2b-c, Extended Data Fig. 7-8). MDA-MB-231 and HCC1806 cells required the pyrimidine synthesis enzyme DHODH to metastasize to all tissues; although, *DHODH* loss in EO771 only reduced metastasis to the liver and kidney/adrenal gland. *GART* was required for all cells to metastasize to every tissue tested, suggesting a possible requirement for purine synthesis to survive in the bloodstream, as this would affect outgrowth in any site. This observation is notable given the relatively high concentrations of the purine salvage precursor hypoxanthine measured in all tissue interstitial fluids and CSF compared to plasma (Fig. 1n, Extended Data Fig. 1d, 2d), and argues that individual nutrient requirements are not sufficient to define the tissues to which these cancer cells can metastasize.

**Fig. 2:**
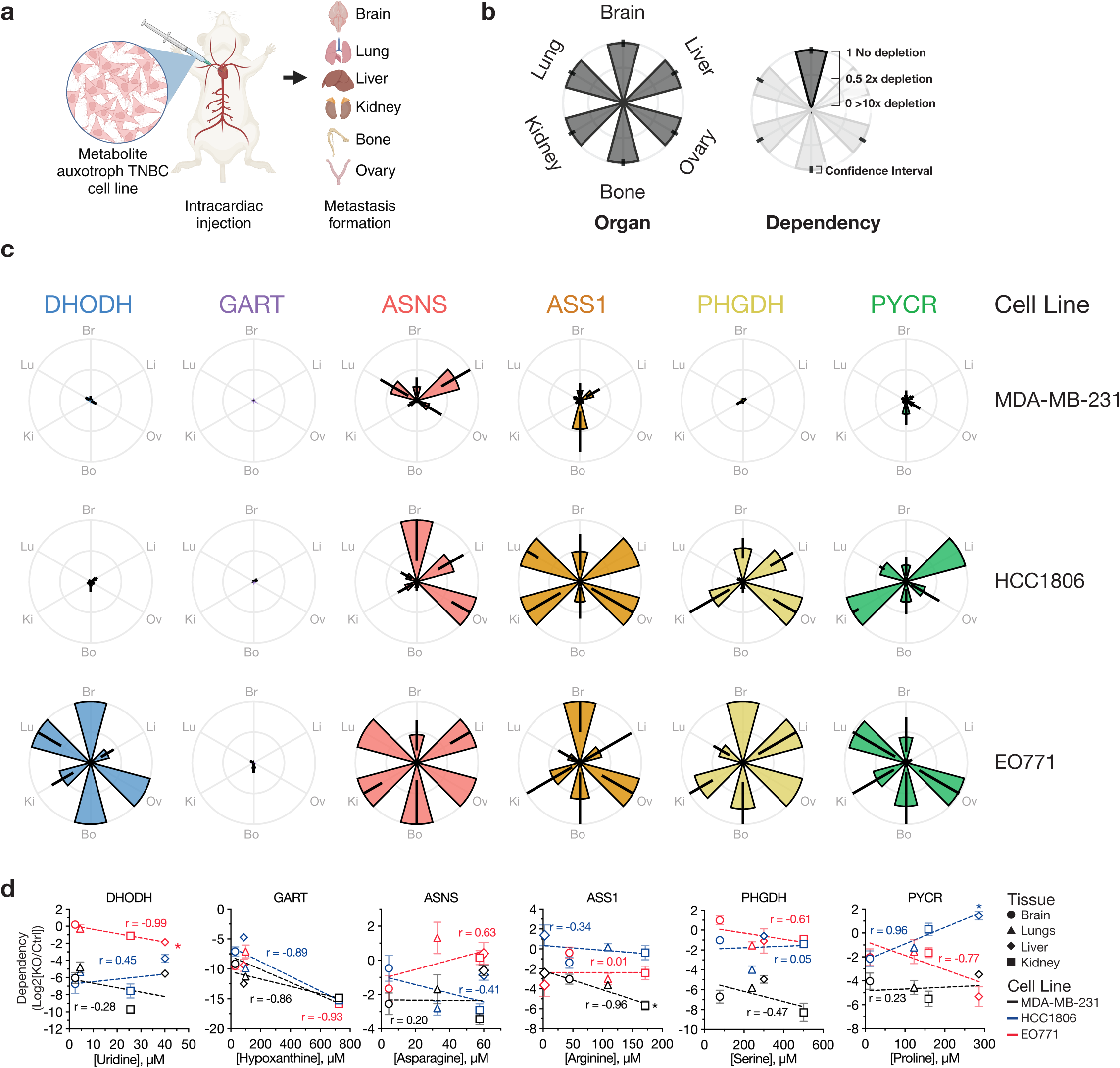
Intracardiac implantation to determine where metabolite auxotrophs can grow as metastases. **a**, Schematic illustrating intracardiac injection of auxotroph and control cells to determine site of metastasis. Both control and auxotrophic cell lines, engineered to express firefly luciferase (Fluc), were injected into the left ventricle of mice, allowing for metastatic spread to various tissues including the brain, liver, lungs, ovaries, bones, and kidneys/adrenal glands. Metastatic colonization was quantified by measuring bioluminescence in harvested tissues at the experimental endpoint. MDA-MB-231-Fluc and HCC1806-Fluc cells were injected into NSG mice, and EO771-Fluc cells were injected into C57BL/6J mice. **b**, Description of petal plots to show the metastatic patterns of metabolite auxotroph cells relative to control cells. Each petal represents a specific tissue, with the length indicating the relative growth of tumors from each metabolite auxotroph cell line compared to control cells. **c**, Petal plots showing the metastatic distributions of different metabolite auxotroph cells relative to control cells. Data are presented as mean ± 95% confidence interval. Raw data used to derive dependency values and number of mice per experimental group are presented in Extended Data Fig. 7-8. **d**, Scatter plots correlating the concentration of relevant metabolites in tissue interstitial fluids with the dependency of each cell lines (black, MDA-MB-231; blue, HCC1806; red, EO771) on the corresponding metabolic genes for metastatic growth in that tissue. The x-axis shows the tissue metabolite concentration relevant to each auxotroph, while the y-axis displays the dependency as a log2 fold change of knockout (KO) compared to control (Ctrl). Symbols denote the metabolite concentration in specific tissues, with brain values derived from CSF measurements. Data represent mean ± SEM, and the raw data used to derive dependency values involve the number of mice per experimental group presented in Extended Data Fig. 7-8. Pearson correlation coefficients (r) and p-values are provided to assess statistical significance (*p < 0.05). PYCR represents PYCR1, 2, and 3 triple KO.

We observed unexpected heterogeneity in the dependency on amino acid synthesis genes to grow in different tissue sites across cell lines (Fig. 2c, Extended Data Fig. 7-8). While HCC1806 cells did not require *ASNS* for metastasis to the brain or ovary, *ASNS* was required for metastasis to the lung, bone, and kidney/adrenal glands. In contrast, *ASNS*-null MDA-MB-231 cells and EO771 cells showed a different pattern of metastasis, with *ASNS* loss slightly affecting brain metastasis of EO771 cells and leading to reduced metastasis of MD-MB-231 cells in all sites except the liver. *ASS1* loss affected metastasis of MD-MB-231 cells to all sites, while *ASS1*-null HCC1806 cells were less able to form tumors in the brain and bone, and *ASS1*-null EO771 cells grew best in brain and ovary. The bone was the site most consistently affected by *ASS1* loss across all three lines, although the reduction in growth from *ASS1*-loss was modest. Consistent with prior studies^15^, MDA-MB-231 cells demonstrated a dependency on *PHGDH* for metastasizing to the brain; however, *PHGDH* was also broadly required for MDA-MB-231 to metastasize to all other tissues examined. Furthermore, HCC1806 only required *PHGDH* for lung and bone metastasis and EO771 demonstrated minimal dependency on *PHGDH* for metastasis to any tissue, only have some effect on lung metastasis. This highlights that despite low serine levels in the brain^19,20^, serine synthesis is not universally required for brain metastasis. Finally, the ability to make proline was required for MDA-MB-231 to metastasize to all tissues, whereas HCC1806 and EO771 displayed variable dependency on proline synthesis to metastasize to the tissues measured. Notably all three cell lines tested showed some dependency on *PYCR1,2,3* for brain metastasis. Collectively, these data indicate that amino acid auxotrophy does not reliably predict tissue-specific metastasis.

Given the heterogeneity in metastatic behavior of metabolite auxotrophs across cell lines, we systematically correlated tissue-specific nutrient levels with metabolic dependencies in order to more formally assess whether nutrient availability could predict tissue-specific metastasis. We hypothesized that a low environmental abundance of a particular nutrient would increase cancer cell dependence on synthesizing that nutrient for survival and growth. For instance, *PHGDH*-null cells would likely exhibit minimal to no colonization defects in serine-rich tissues but would struggle to establish metastases in serine-poor tissues. While our data confirmed a positive correlation between proline concentration and *PYCR1,2,3* dependency in HCC1806 cells, a similar correlation was not found for the other auxotrophic nutrients (Fig. 2d). In fact, we found a significant negative correlation between *ASS1* and arginine levels in MDA-MB-231 cells and for *DHODH* and pyrimidine levels in EO771 cells. Furthermore, correlating all measured metabolites with metastatic potential revealed only a few instances where single nutrient levels correlated with dependency on any of the metabolic synthesis genes tested, and these correlations were typically unrelated to the metabolite auxotrophy being tested (Extended Data Fig. 9). These findings challenge the assumption that levels of individual nutrients in tissues directly predict metabolic pathway dependencies to metastasize and grow in a tissue.

### Assessing gene-nutrient dependencies to form tumors when cells are directly implanted into the brain or MFP

Given the possibility that nucleotide synthesis (and thus *DHODH* or *GART* expression) is required for survival in circulation, and the unexpected lack of correlation between tissue nutrient availability and the dependency on metabolic genes for metastasis, we sought to further test the ability of tissue nutrient levels to predict whether specific auxotrophs can form tumors when directly implanted into a tissue site. Specifically, we used direct implantation of MDA-MB-231 or HCC1806 cells into the brain or MFP of mice to assess tissue colonization separate from other factors associated with metastasis, including survival in circulation (Fig. 3a). We chose the brain and MFP for these studies since the availability of many nutrients, especially amino acids, is overall lower in the brain microenvironment compared to circulation^19,20^ (Fig. 1d, l-n, Extended Data Fig. 1a-j, 2c-d) and prior work suggested in some cases cancer cells can grow best in the primary site nutrient environment^42^. For these experiments, each auxotroph was engineered to express Gaussia luciferase (Gluc)^43^, enabling the monitoring of tumor burden by quantification of blood luminescence. We also injected each EO771 auxotroph or control cell line into the brain to specifically compare brain tumor metabolic gene dependency across all three cell lines (Fig. 3b-d, Extended Data Fig. 10-11).

**Fig. 3:**
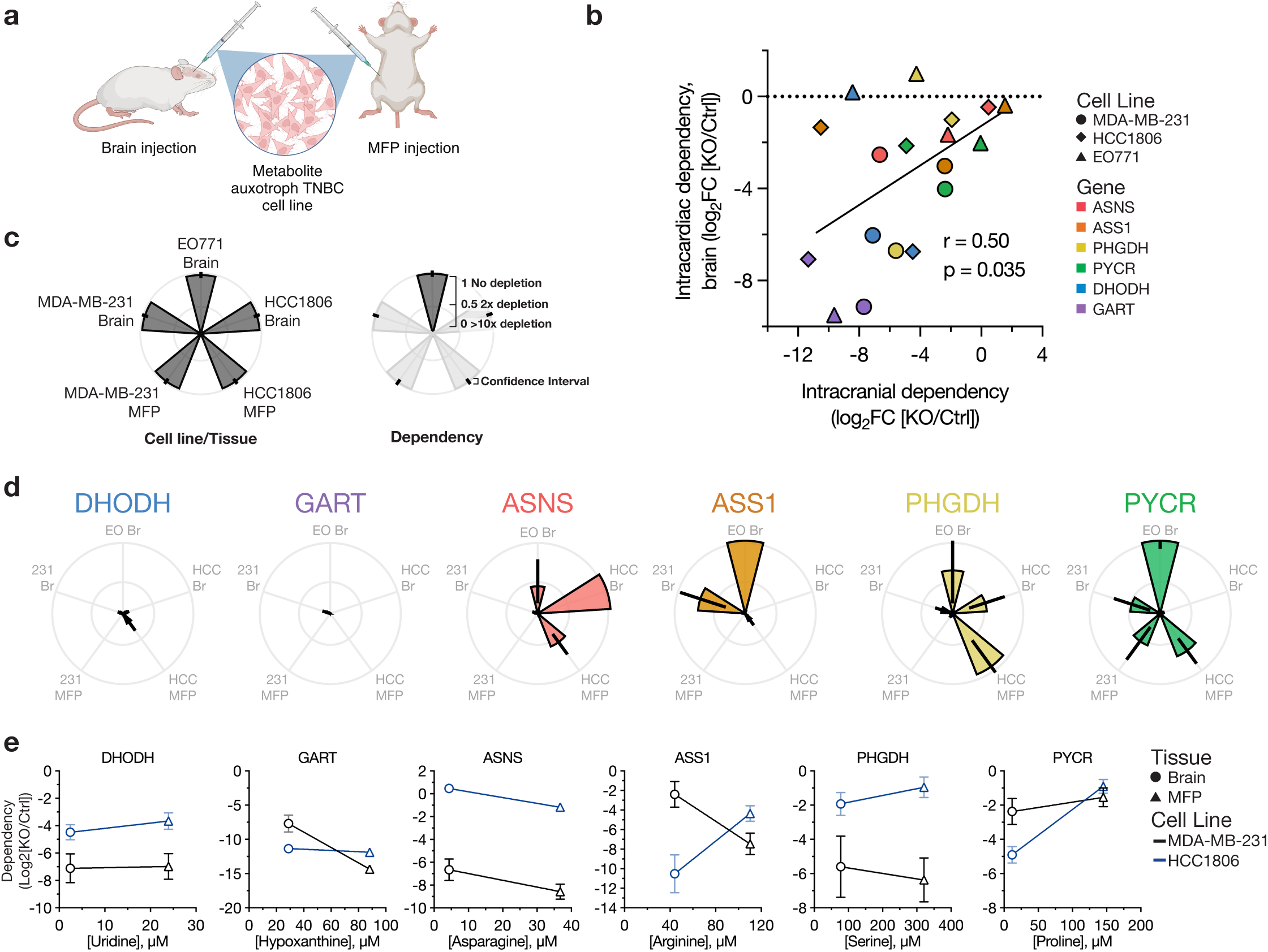
Assessing metabolic dependencies of brain and MFP tumors. **a**, Schematic illustrating use of direct implantation of auxotrophs and control cells into brain and mammary fat pat (MFP) to assess metabolic dependencies. Both control and auxotrophic cell lines, engineered to express Gaussia luciferase (Gluc), were injected into the brain or MFP of mice, then tumor growth monitored over time via blood luminescence. MDA-MB-231-Gluc and HCC1806-Gluc cells were injected into NOD-SCID-gamma mice, and EO771-Gluc cells were injected into C57BL/6J mice. **b**, Scatter plot correlating the dependency of cell lines on specific metabolic genes for metastasis to the brain based on route of cell delivery into mice. The x- and y-axes show the average dependency values as a log2 fold change of knockout (KO) compared to control (Ctrl) cells introduced into mice via intracranial or intracardiac injections, respectively. The number of mice per experimental group are indicated in Extended Data Fig. 7-8 and Extended Data Fig. 10. r and p-values were determined by Pearson correlation. **c**, Description of petal plots to show the tumor growth patterns of metabolite auxotroph cells. Each petal represents a specific cell line and tumor, with the length indicating the relative growth of tumors from each metabolite auxotroph cell line compared to control cells. **d**, Petal plots showing tumor growth of different metabolite auxotroph cells relative to control cells. Data are presented as mean ± 95% confidence interval. Raw data used to derive dependency values and number of mice per experimental group are presented in Extended Data Fig. 10. **e**, Scatter plots illustrating the relationship between the concentrations of relevant metabolites in MFP interstitial fluid and CSF, representing brain metabolite levels, and the dependency of cell lines (black, MDA-MB-231; blue, HCC1806) on the corresponding metabolic genes for metastatic growth in each tissue site. The x-axis denotes the tissue metabolite concentration relevant for each metabolite auxotroph cell line, while the y-axis indicates the dependency as log2 fold change of each indicated KO relative to Ctrl. Symbols denote the metabolite concentration in specific tissues, with brain values derived from CSF measurements. Data represent mean ± SEM, and the raw data used to derive dependency values and number of mice per experimental group are presented in Extended Data Fig. 10. PYCR or PYCR KO represents PYCR1, 2, and 3 triple KO.

We first compared how the route of cell delivery to the brain influenced gene dependency for tumor growth in that site. Our analysis revealed a significant correlation between gene dependencies observed with intracranial and intracardiac injections (Fig. 3b, Extended Data Fig. 11g), indicating that the overall patterns of gene dependency are independent of the method of cell delivery to the brain. However, we found that dependencies were generally stronger when cells were directly implanted via intracranial injection. Additionally, we observed some notable outliers where the route of injection affected gene dependency. For instance, HCC1806 cells demonstrated a strong dependency on *ASS1* when injected intracranially, but this dependency was reduced following intracardiac injection. EO771 cells lacking *DHODH* or *PHGDH* were able to form metastatic brain tumors when injected in the heart but not when directly implanted in the brain.

In line with the intracardiac injection tumor formation data, dependency on nucleotide synthesis was invariably required for metastatic growth in the brain, as loss of either *DHODH* or *GART* reduced brain tumor growth across all three cell lines tested and extended mouse survival (Fig. 3d, Extended Data Fig. 10, 11a-b). These effects were also observed at the primary tumor site for MDA-MB-231 and HCC1806, again mirroring intracardiac injection tumor formation data, and showing a broad requirement for de novo nucleotide synthesis to enable growth of these cells in multiple tissues despite high levels of nucleotides measured in tissue IF (Fig. 1m-n, Extended Data Fig. 1d, 2d). These experiments also argue against an inability to survive in the circulation to explain reduced growth of nucleotide auxotrophs in different tissues based on intracardiac injections.

Consistent with the intracardiac injection tumor formation experiments, we again observed heterogeneity in dependency on amino acid synthesis genes to grow in tissues across cell lines (Fig. 3d, Extended Data Fig. 10, 11c-f). For instance, MDA-MB-231 required *ASNS* for growth in both the brain and MFP, whereas EO771 and HCC1806 showed intermediate or no dependency on *ASNS* to form brain tumors, respectively. In contrast, HCC1806 displayed a strong dependency on *ASS1* in both brain and MFP tumors, while MDA-MB-231 cells required *ASS1* exclusively in the MFP and EO771 did not require *ASS1* to grow in the brain. We again observed that MDA-MB-231 cells exhibited a dependency on *PHGDH* for brain tumor growth, with knockout of this gene significantly extending survival of mice with *PHGDH*-null MDA-MB-231 brain tumors, consistent with published work^15^. However, this dependency on *PHGDH* in MDA-MB-231 cells was also evident in MFP tumors, suggesting that the requirement for *PHGDH* cannot be solely attributed to tissue serine levels as serine was measured to be higher in MFP (Fig. 1l, Extended Data Fig. 1a). Consistent with intracardiac injection tumor formation results, HCC1806 and EO771 displayed a variable requirement for *PHGDH* to grow in the brain (Fig. 3d, Extended Data Fig. 10, 11e). HCC1806 *PHGDH* knockout cells grew slower in the brain compared to control, whereas EO771 displayed minimal reliance on *PHGDH* for brain tumor growth with negligible impact on mouse survival. Both MDA-MB-231 and HCC1806 cells showed some dependence on *PYCR1,2,3* for tumor growth in brain. Additionally, HCC1806 showed a reduced dependence on *PYCR1,2,3* for MFP tumor growth compared to the brain, suggesting that proline synthesis may be a more brain-specific dependency; however, EO771 cells were not dependent on *PYCR1,2,3* for brain tumor growth. From these experiments, a consistent link between tissue nutrient availability and tumor growth was not observed (Fig. 3e). We noted a trend in the case of *ASS1*, *PHGDH*, and *PYCR1,2,3* dependency in HCC1806 cells, but this trend was not evident in MDA-MB-231 cells. Collectively, these data reinforce that dependency on single nutrients or single nutrient levels is not a reliable general predictor of metabolic gene dependency for tumor growth in different sites. Furthermore, these data suggest a more complex interplay between multiple nutrients or environmental factors and cell-intrinsic properties to enable the growth of cancer cells as metastases in different tissues.

### Comparative analysis of metabolite synthesis activity and nutrient dependency in primary tumors versus brain metastases

To further study the relationship between nutrient levels in tissues and the ability to synthesize specific nutrients to allow tumor growth in a given tissue, we traced the fate of ^13^C-labeled glucose in MDA-MB-231-derived brain or MFP tumors in conscious unrestrained mice (Fig. 4a, Extended Data Fig. 12a). We confirmed that the amount of glucose label in the plasma reached steady-state over the infusion period (Fig. 4b). We then quantified the incorporation of ^13^C-label into metabolites synthesized downstream of glucose in rapidly isolated tumor-bearing and adjacent non-tumor-bearing tissue in order to understand how tumors acquire different nutrients. Labeling of lactate and TCA cycle metabolites was higher in MFP tumors, brain, and brain tumors compared to MFP (Extended Data Fig. 12b-c). The similar levels of ^13^C label in non-tumor brain tissue and tumors growing in both sites likely reflect the high glucose utilization in brain tissue^44^ and tumors^45^. Of note, we observed more labeling of asparagine, glycine, serine, and proline in brain tumors compared to MFP tumors, suggesting higher rates of synthesis of these amino acids in brain tumors (Fig. 4c-f, Extended Data Fig. 12d-e). Despite these labeling differences, MDA-MB-231 cells null for *ASNS*, *PHGDH*, or *PYCR1,2,3* were similarly defective in their ability to grow as tumors when implanted into either the MFP or the brain (Fig. 3d). This indicates that heightened amino acid synthesis activity in the brain does not necessarily reflect an increased dependency on these pathways for tumor growth in brain, despite the lower availability of most amino acids in the CSF (Fig. 1l, Extended Data Fig. 1a, 2c). These findings further suggest that the relatively lower amino acid synthesis activity observed in MFP tumors is sufficient to sustain tumor growth in that site.

**Fig. 4:**
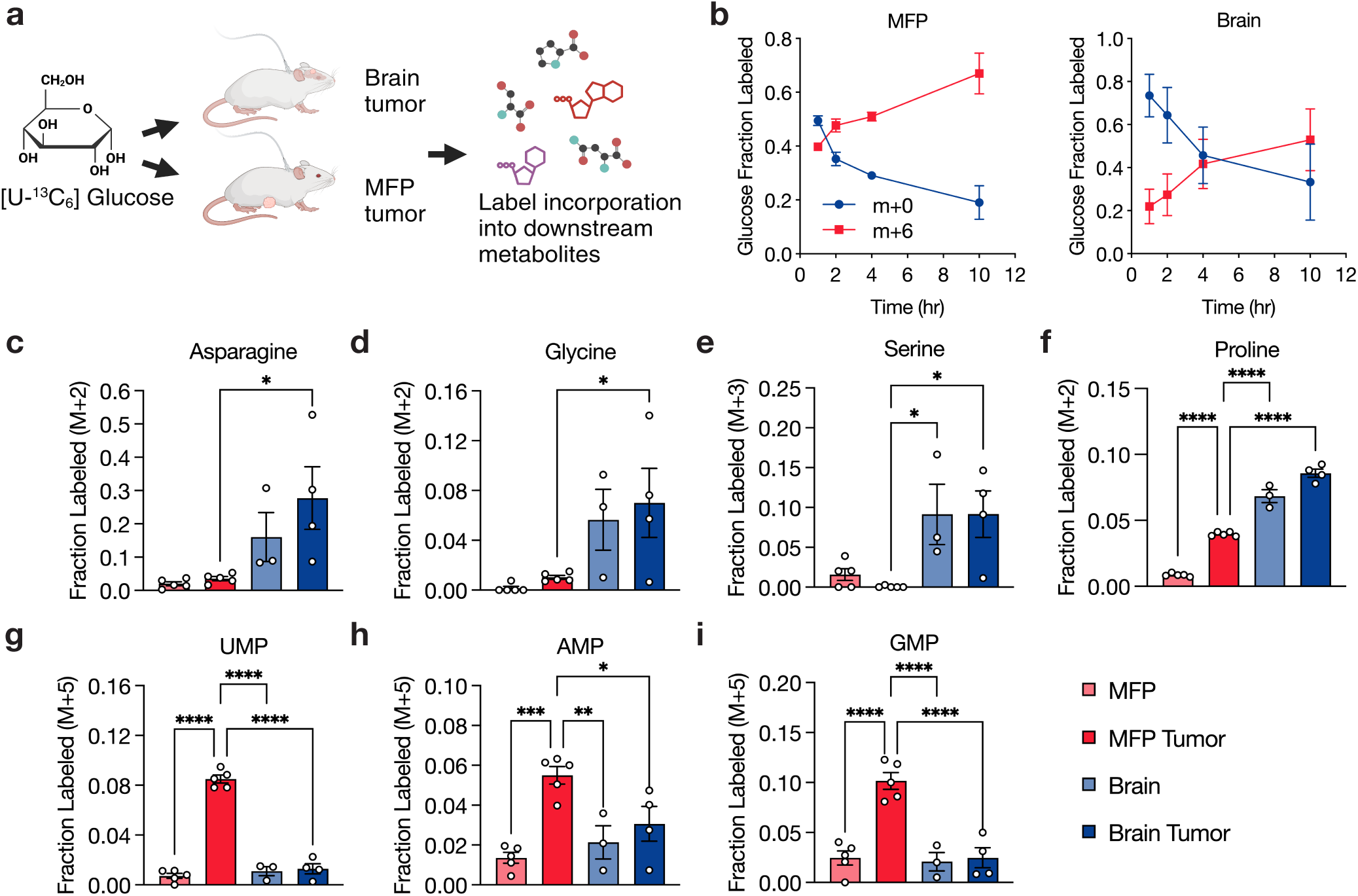
Assessment of metabolite fate in primary and brain metastatic breast cancers. **a**, Schematic showing use of ^13^C-glucose (m+6) to assess metabolite fate in female NSG mice bearing MDA-MB-231 cell-derived tumors located either in the mammary fat pad (MFP) or brain. **b**, Fractional labeling of glucose in the plasma (m+0 or m+6) following infusion with [U-^13^C]-glucose at a rate of 0.4 mg/min for 10 hours in female NSG mice harboring either MFP or brain MDA-MB-231 tumors. Data points represent mean ± SEM for n = 5 (MFP tumor) or n = 4 (brain tumor) biological replicates. **c-i**, Fractional labeling of specific metabolites determined by LC/MS in cancerous tissues (MDA-MB-231 tumors in the brain and MFP) and corresponding noncancerous tissues (brain and MFP) from NSG mice infused with [U-^13^C]-glucose. Data points represent mean ± SEM for n = 5 (MFP tumor, MFP), 3 (noncancerous brain), and 4 (brain tumor) biological replicates. Statistical analysis was performed using an ordinary one-way ANOVA with Holm-Sidak’s multiple comparisons test (*p < 0.05, **p < 0.01, ***p < 0.001, ****p < 0.0001).

In analyzing nucleotide synthesis, we found that MDA-MB-231 MFP tumors synthesized more pyrimidine and purine nucleotides from glucose compared to normal MFP tissue (Fig. 4g-i, Extended Data Fig. 13), a finding consistent with the observation that tumors undergo elevated rates of nucleotide synthesis than non-transformed tissues^46^. Notably, nucleotide synthesis was lower in both brain and brain tumors compared to MFP tumors, and labeling in these tissues was similar to normal MFP. This may reflect faster proliferation of cancer cells in the MFP relative to the brain. This trend also suggests the brain may rely more on salvage than de novo synthesis to maintain sufficient nucleotide levels^47^, which is consistent with work showing lower purine synthesis activity in the brain compared to other tissues^48,49^. To determine whether differences in nucleotide availability between MFP and the brain could explain this phenotype, we considered nucleotide levels in both MFP IF and CSF (Fig. 1m-n, Extended Data Fig. 1d). We observed that, with the exception of IMP, all the nucleotides or nucleobases measured in MFP IF versus CSF were lower in CSF. Nevertheless, despite tissue site differences in labeling downstream of nucleotide synthesis, MDA-MB-231 cells were still dependent on *DHODH* and *GART* for growth in both brain and MFP tumors (Fig. 3d). These data argue that neither the levels of individual metabolites nor the activity of individual pathways in tumors can always predict where cancers can metastasize. Rather, it implies that tumor cells differentially leverage existing nutrient resources or engage alternative pathways within a tissue microenvironment to meet their metabolic demands for tumor growth.

### Analyzing metastatic potential through gene expression and CRISPR dependency

To further our understanding of metabolic requirements and metastasis, we analyzed public data from the Broad Institute Dependency Map portal and correlated metastatic potential with gene expression and CRISPR dependency across breast cancer cell lines for the metabolic genes studied here. Our analysis revealed no significant correlation between tissue specific metastatic potential and expression of the metabolic genes considered in this study (Extended Data Fig. 14). However, significant correlations were observed between CRISPR dependencies for *DHODH* and *GART* and metastatic potential involving several tissues, particularly the lungs (Fig. 5a, Extended Data Fig. 15). This is notable given that we found *GART* and *DHODH* to be essential in both MDA-MB-231 and HCC1806 cells for metastasis to all tissues (Fig. 2c). For the other genes considered, no significant correlations were found between CRISPR dependency and metastatic potential (Fig. 5a, Extended Data Fig. 15). These findings suggest that CRISPR dependencies of metabolic genes may provide more reliable insights into tissue-specific metastatic potential than gene expression or levels of tissue metabolites in select cases.

**Fig. 5:**
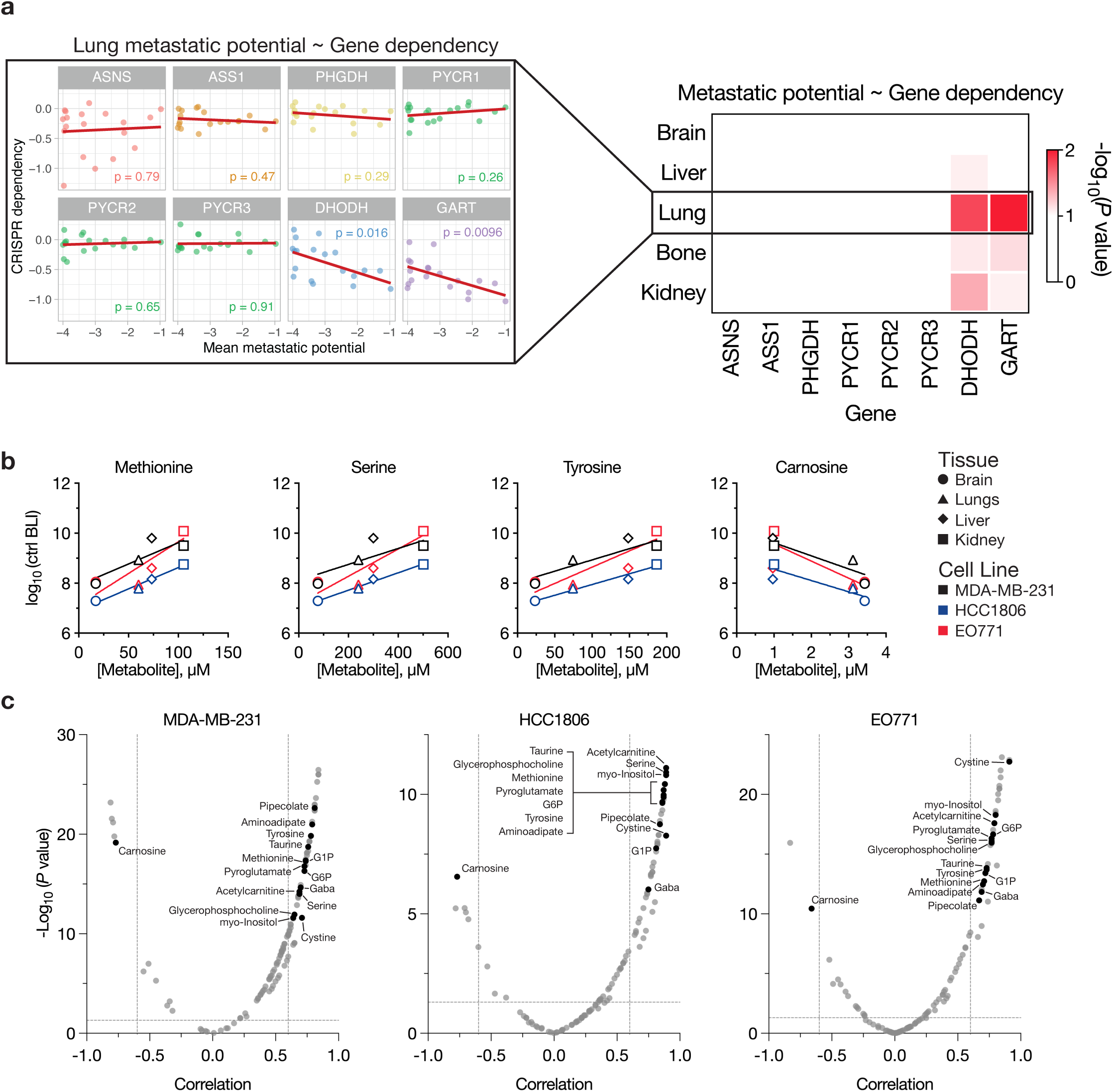
Correlating metabolic gene dependencies and metabolite levels with tissue-specific metastatic potential. **a**, Left, scatter plots correlating the metastatic potential of breast cancer cell lines to the lung with in vitro CRISPR dependencies of the indicated genes. Each dot represents a cell line, and p-values derived from Pearson correlations are indicated within each plot. Right, heat map showing the −log_10_(p-values) from correlating metastatic potential of breast cancer cell lines to the indicated tissues with CRISPR dependency of the indicated genes. Data sourced from the Dependency Map portal. **b**, Scatter plots correlating the concentration of relevant metabolites in tissue interstitial fluids with the metastatic potential of control (ctrl) cells (black, MDA-MB-231; blue, HCC1806; red, EO771) following intracardiac injection. Symbols denote the metabolite concentration measured in specific tissues, with brain values derived from CSF measurements. Data represent mean ± SEM, and the raw data used to derive metastatic potential values are presented in Extended Data Fig. 7-8. For all graphs, |Pearson correlation, r| > 0.6 and p-values < 0.0001. **c**, Volcano plots depicting the Pearson correlation values and p-values for metabolites correlated with metastatic potential of control cell lines following intracardiac injection. Black circles indicate metabolites that are significantly correlated in all three cell lines tested in this study. Cutoffs of |Pearson correlation, r| > 0.6 and p-values < 0.05 were used to select significantly correlated metabolites. G6P: Glucose-6-phosphate; G1P: Glucose-1-phosphate.

### Multiple metabolites influence tissue-specific metastatic potential

From our analyses, we noticed that the brain microenvironment is the most depleted tissue environment for many nutrients (Fig. 1d, l-n, Extended Data Fig. 1a-j, 2c-d), and it is also the tissue with the slowest MDA-MB-231 and HCC1806 metastatic tumor growth (Extended Data Fig. 6b). Therefore, we investigated whether levels of any nutrients could predict metastatic outgrowth in control cell lines at baseline. To this end, we correlated the metastatic potential of the control cell lines from the intracardiac injection experiments with levels of metabolites measured in the brain, liver, lung, and kidney. This analysis revealed that several metabolites positively correlate with metastatic potential (Fig. 5b-c). Notably, 15 metabolites consistently exhibited significant correlations across all three cell lines, pointing to their potential importance in predicting metastatic potential. This included several amino acids, including serine, methionine and tyrosine. In contrast, some metabolites such as carnosine negatively correlated with metastatic potential, indicating that their presence might inhibit tumor growth in certain tissues. Together, these findings suggest that the ability of cancer cells to metastasize to specific tissues is likely influenced by the combined levels of multiple metabolites within tissues, making predictions based on single metabolites insufficient. Therefore, a deeper understanding of the collective influence of nutrient levels across different tissues could provide more accurate insights into the factors that drive metastatic potential and tumor growth.

## Discussion

A consistent correlation between tissue-specific single nutrient levels and the dependency on specific metabolic genes was not observed across three breast cancer cell lines. While extensive data supports the notion that nutrient availability influences metabolic behavior^5,50^, these findings suggest that metabolic dependencies in cancer are not solely determined by the availability of metabolites synthesized downstream of a given metabolic enzyme. Rather, the data indicate that metabolic dependencies and tissue preference of metastases are shaped by a complex interplay between multiple nutrients in tissues and cell-intrinsic factors that influence tumor metabolic requirements. For example, serine, which is essential for protein, lipid, and nucleotide synthesis^51^, was confirmed to be lower in the brain microenvironment^19,20^. Nevertheless, we observed that serine synthesis via PHGDH is not universally required across cell lines for brain tumor growth or brain metastasis. One possibility is that different cell lines can differentially salvage products of serine synthesis, influencing their dependence on de novo synthesis. Additionally, cancer cell lines do not uniformly respond to serine starvation, even within the same primary tumor type, indicating inherent differences in serine metabolism across cell lines^52^. Furthermore, serine synthesis activity can be influenced by the ability of cells to respond to other environmental nutrient levels; this implies a metabolic budget exists whereby cells are wired in a manner that addicts them to uptake of specific nutrients to enable use of limiting metabolic resources to make what is not present^52^. This may underlie why cells retain a preference to grow in some tissue nutrient conditions even when repeatedly exposed to a different tissue nutrient environment^42^. Moreover, the dependency on metabolic genes such as *PHGDH* may also fluctuate throughout different stages of the metastatic cascade. For instance, low *PHGDH* expression may drive early-stage metastasis in breast cancer, whereas re-expression occurs in established metastases^53^. This suggests that the reliance of certain cancer cells on serine synthesis or other metabolic pathways is dictated by a complex interaction of cell-extrinsic factors within the tumor microenvironment and cell-intrinsic metabolic demands, complicating the prediction and targeting of metabolic vulnerabilities based on a small number of variables.

Our quantification of metabolites available across tissues challenge the conventional view that tissue microenvironments are generally nutrient-deprived. Supporting this, tumor interstitial fluid is not depleted of many nutrients^24,54^. Notably, tumor interstitial fluid is similar to normal kidney interstitial fluid in human patients with renal cell carcinoma, suggesting that tumors adapt to existing environmental conditions rather than reshaping them to suit their needs^24^. This adaptation is particularly critical in the brain, where the microenvironment is more nutrient-restricted, largely due to the selective permeability of the blood-brain barrier^55^. Despite lower levels of many metabolites in the brain microenvironment, we did not find a universal dependency on key amino acid synthesis pathways, underscoring that differences in cell-intrinsic metabolic traits and the ability to salvage nutrients from the microenvironment likely drive these varied dependencies.

Previous studies have shown that heightened tumor metabolic synthesis activity, as measured by labeled glucose incorporation, often correlates with dependency on the synthesis pathway, such as for serine or lipid synthesis in the brain^12,13,15^. However, our results suggest that this correlation is not always the case. This discrepancy may stem from the fact that we conducted bulk tumor measurements, which may mask tumor cell-specific metabolic behaviors. Alternatively, genetic knockouts may induce metabolic adaptations, such as nutrient salvage, allowing tumor cells to utilize available nutrients in the tissue microenvironment and reduce dependency on the synthesis pathway. An example of this adaptation was observed in pancreatic cancer cells, which increase uridine salvage from culture medium in response to the loss of *DHODH* and pyrimidine synthesis activity^56^.

Despite the heterogeneity in metabolic dependencies we observed in auxotrophs, we found a strong dependency on *GART* for breast cancer to grow in multiple tissues. This robust dependency on *GART* underscores the importance of purine synthesis in this cancer and is corroborated by evidence that breast cancer can be treated with antifolates^57^. Notably, we did not observe elevated purine synthesis in brain compared to MFP tumors, which is consistent with two recent studies showing lower purine synthesis activity in the brain compared to other tissues^48,49^. While those data were interpreted as a reliance of brain tissue on nucleoside or nucleobase salvage^47^, the results here argues that synthesis remains important for tumor growth and that neither the amount of de novo synthesis activity nor the levels of salvage precursors are necessarily predictive of relative pathway dependency.

*ASS1* downregulation is prevalent in various cancers, including breast cancer^38^, leading to arginine deprivation as a proposed therapeutic strategy^58^. However, the effectiveness of arginine depletion has been inconsistent in clinical studies^59^. Given that *ASS1* dependency was not found to be well-predicted by arginine availability, additional factors are necessary to select which patients are most likely to responds to arginine depleting therapies.

Our use of different models to explore gene-nutrient interactions highlights how methodological approaches can lead to different interpretations of metabolic dependencies. For example, while *DHODH* was found to be essential for MDA-MB-231 and HCC1806 cells across all experimental approaches, *DHODH* was required by EO771 cells for brain and MFP tumor growth but was minimally essential for metastasis when introduced into mice via intracardiac injections. Differences in metabolic dependencies between intracardiac and intracranial injections likely reflect the distinct microenvironments and stages of metastasis encountered with each method. Despite these differences, each method offers insights into understanding the metabolic requirements of cancer cells for metastatic tumor growth.

This study only examined breast cancer, and whether the relationship between single nutrient levels and metabolic requirements for metastasis extends to other cancer types will require future studies. Moreover, tumors arising from the clonal knockout cells that are necessary to create auxotrophs may not capture intratumor metabolic heterogeneity that could contribute to nutrient sharing among cancer cells within a tumor^53,60^. Despite these limitations, this study provides insights into the metabolic dependencies of TNBC cells and their relationship with nutrient availability across different tissues. That is, the interplay between levels of multiple microenvironmental nutrients and cell-intrinsic metabolic wiring should be considered in order to better understand the metabolic requirements of metastasis.

## Supporting information

Supplementary Table 1 Metabolites measured in this study

Supplementary Table 2 Metabolite concentrations tissue IF, CSF, and plasma from NSG and C57BL/6J mice

Supplementary Table 3 Metastatic potential of auxotroph cell lines following intracardiac injections

Supplementary Table 4 Tumor growth of auxotroph cell lines following intracranial or MFP injections

Supplementary Table 5 Correlations of metastatic potential with tissue metabolite concentrations

Supplementary Table 6 Primer sequences used in this study

## Acknowledgements

We thank members of the Vander Heiden, Jain, and Church laboratories for helpful discussions and the Koch Institute’s Robert A. Swanson (1969) Biotechnology Center for technical support, specifically the Preclinical Imaging & Testing, Flow Cytometry, Barbara K. Ostrom (1978) Integrated Genomics and Bioinformatics, and Hope Babette Tang (1983) Histology facilities. We also thank the MIT Division of Comparative Medicine staff and the Massachusetts General Hospital Center for Comparative Medicine staff for help with colony maintenance and animal care, and Christopher S. Nabel for comments on the manuscript. This work was supported in part by R01CA259253 to R.K.J. and M.G.V.H. as well as the Koch Institute Cancer Center Support Grant P30CA014051 and the Koch Institute/Dana-Farber Harvard Cancer Center Bridge project. K.L.A. was supported by the National Science Foundation (DGE-1122374) and National Institutes of Health (NIH) (F31CA271787, T32GM007287). S.Subudhi was supported by the MGH ECoR FMD fellowship grant (2022A018897). R.F. was supported by the Knut and Alice Wallenberg Foundation (KAW 2019.0581). S.E.H. acknowledges support from the Marietta Blau-Grant by the Austrian Agency for Education and Internationalisation. S.Sivanand acknowledges support from the Damon Runyon Cancer Research Foundation (DRG-2367-19). R.F., L.M.R., and G.M.C. were supported by the Aging and Longevity-Related Research Fund and EGL Charitable Foundation. A.A. received support as a Howard Hughes Medical Institute (HHMI) Medical Research Fellow. X.J. acknowledges support from Key R&D Program of Zhejiang (2024SSYS0034). R.K.J. is supported by grants from the NIH (U01CA224348; R01CA208205; R01NS118929; U01CA261842), the Ludwig Cancer Center at Harvard, Nile Albright Research Foundation, National Foundation for Cancer Research, and Jane’s Trust Foundation. M.G.V.H. acknowledges support from the MIT Center for Precision Cancer Medicine, the Ludwig Center at MIT, and the NIH (R35CA242379).

## Author Contributions

Conceptualization: K.L.A., S.Subudhi, R.F., A.A., X.J., R.K.J., M.G.V.H.; Methodology: K.L.A., S.Subudhi, R.F., Y.G., A.A., N.H., V.S., F.J.S.R., X.J., R.K.J., M.G.V.H.; Investigation: K.L.A., S.Subudhi, R.F., Y.G., S.C.S., M.B.M., S.E.H., A.S.K., D.L.R., M.W., J.A.H., S.Sivanand, L.M.R., M.D., A.A., N.H., A.S.Z., F.G., A.M.B., M.W., T.K., G.B.F., B.T.D.; Resources: G.M.C., R.K.J., M.G.V.H.; Writing – Original Draft: K.L.A., S.Subudhi, R.F., R.K.J., M.G.V.H.; Writing – Review & Editing: All authors; Supervision: V.S., F.J.S.R., X.J., G.M.C., R.K.J., M.G.V.H.; Funding Acquisition: G.M.C., R.K.J., M.G.V.H. Y.G., S.C.S., M.B.M., contributed equally to this study.

## Lead contact

Further information and requests for resources and reagents should be directed to and will be fulfilled by Matthew G. Vander Heiden, George M. Church, or Rakesh K. Jain.

## Methods

### Cell lines and culture

Human cell lines used in this study were obtained from ATCC (HCC1806: CRL-2335, from female patient; MDA-MB-231: HTB-26, from female patient). Murine EO771 breast cancer cells were obtained from CH3 BioSystems (94A001, from C57BL/6 mice). Lines were regularly tested for mycoplasma contamination using the MycoAlert PLUS Mycoplasma Detection Kit (Lonza BioSciences, LT07-710). All cells were cultured in a Heracell humidified incubator (Thermo Fisher Scientific) at 37 °C and 5% CO_2_. Cell lines were routinely maintained in RPMI-1640 (Corning Life Sciences, 10-040-CV) supplemented with 10% heat inactivated fetal bovine serum (Gibco, 10437-028), and for cell culture experiments, 10% dialyzed fetal bovine serum (Gibco, 26400-044) was supplemented.

RPMI-1640 medium lacking serine or arginine was made using the method outlined in (Muir et al. 2017^61^). Briefly, enough of all amino acids and sodium phosphate dibasic (Sigma-Aldrich, S5136) of RPMI-1640 media except for serine or arginine were weighed out to make 25 L of media, then the resulting powders were homogenized using an electric blade coffee grinder (Hamilton Beach, 80365) that had been washed with methanol then water. The resulting powders were resuspended in water along with sodium bicarbonate (Sigma-Aldrich, S5761) and RPMI-1640 medium w/o amino acids, sodium phosphate (US Biological, R8999-04A) to make RPMI-1640 media lacking serine or arginine. Serine or arginine was dissolved in water to make 1000-fold concentrated stock solutions, then added back to RPMI-1640 media lacking serine or arginine to achieve 286 µM serine or 1.15 mM arginine, respectively.

### Animal studies

All experiments conducted in this study were approved by the MIT Committee on Animal Care (CAC) or MGH Institutional Animal Care and Use Committee (IACUC). Female C57BL/6J and female NOD-SCID-gamma (NSG) mice at 6-10 weeks of age were used in this study. All animals were housed at ambient temperature and humidity (18–23 °C, 40–60% humidity) with a 12 hr light and 12 hr dark cycle and co-housed with littermates with ad libitum access to water. All experimental groups were age-matched, numbered and randomly assigned based on treatment. Data were collected from distinct animals, where n represents biologically independent samples. No statistical methods were used to predetermine sample size. Investigators were not blinded to allocation during experiments and outcome assessment.

### Isolation of tissue interstitial fluid, cerebrospinal fluid, and plasma

Interstitial fluid was collected from mouse tissues using an adapted protocol based on prior published work^54^. Female NSG mice on ad libitum diet aged 6-9 weeks were used for all tissue interstitial fluid isolations. Organs from five mice were combined per interstitial fluid sample, and each pooled sample was treated as an individual data point for the LC/MS measurements and analysis. All mice were euthanized at the same time of day to account for any effects of time of day on metabolism. Tissues were kept on ice throughout the harvest and when ready to pool, were briefly rinsed in ice-cold saline (Azer Scientific, 16005), excess liquid removed by blotting on filter paper (VWR, 28298–020), then placed in a 50 mL conical vial lined with a 20 μm nylon mesh filter (Spectrum Labs, 148134) and 0.25 μL of 0.5 M EDTA, pH 8.0 at the bottom. The tissues were centrifuged at 400 x g for 10 min at 4 °C. The flow through was collected and centrifuged again at 400 x g for 10 min at 4 °C prior to flash freezing and storage in −80 °C until further analysis. Prior to tissue harvest, plasma from each live mouse was collected by facial cheek bleed into EDTA-coated tubes (Sarstedt, 41.1395.105), then centrifuged at 800 x g for 10 min at 4 °C. Supernatant containing plasma was flash frozen and stored at −80 °C until further analysis. An equal volume of each plasma sample was pooled from each mouse cohort prior to analysis by LC/MS.

Cerebrospinal fluid (CSF) was isolated from mouse brain as previously described^62^. Briefly, female NSG or female C57BL/6J mice were anesthetized with ketamine (90 mg/kg) and xylazine (9 mg/kg) via intramuscular injection and CSF collected by gently inserting a sharpened capillary (inner diameter 0.75 mm, outer diameter 1.0 mm) in the cisterna magna. Collected fluid was visually monitored for blood contamination. CSF was centrifuged at 800 x g for 10 min at 4 °C, and the supernatant was flash frozen and stored at −80 °C until further analysis. Blood was collected from the same animal via cardiac puncture and immediately placed in EDTA-tubes (Sarstedt, 41.1395.105), then centrifuged at 800 x g for 10 min at 4 °C. Plasma was flash frozen in liquid nitrogen and stored at −80 °C until further analysis.

### MFP and brain tumor model generation

MDA-MB-231, HCC1806, and EO771 cell lines were transduced with a lentiviral vector expressing Gaussia luciferase (Gluc) and GFP separated by an internal ribosomal entry site^43^, and the top 10% of GFP+ cells were isolated by FACS. To generate MFP tumors, mice were first anesthetized with ketamine (90 mg/kg) and xylazine (9 mg/kg) via intraperitoneal injection, and 1E5 MDA-MB-231-Gluc or HCC1806-Gluc cells were injected into the mammary fat pad in 30 or 50 µL volume of PBS. To generate brain tumors, mice were first anesthetized with ketamine (90 mg/kg) and xylazine (9 mg/kg) via intramuscular injection. Then, 1E5 MDA-MB-231-Gluc, 1E5 HCC1806-Gluc, or 5E4 EO771-Gluc cells were diluted in 1 µL PBS and stereotactically injected into the left frontal lobe of the mouse brain. Tumor volumes were assessed 1-2 times per week by mixing 7 µL tail vein blood with 7 µL 0.5 mM EDTA (pH 8.0) and quantifying luminescence using Promega GloMax Plate Reader (Promega) and the substrate coelenterazine (NanoLight Technology, 303).

### Intracardiac injection and metastasis quantification

MDA-MB-231, HCC1806, and EO771 cell lines were transduced with a lentiviral vector expressing Firefly luciferase (Fluc) and GFP^12^, and infected cells were sorted by FACS using a fixed gate for GFP. For intracardiac injections, 1E5 cancer cells were suspended in 100 µL PBS and injected into the left ventricle of mice anesthetized with inhaled isoflurane using ultrasound guidance^63^. In vivo metastasis progression was monitored via real-time bioluminescence imaging (BLI) using the IVIS Spectrum Imaging System (PerkinElmer). Mice were anesthetized with inhaled isoflurane, injected intraperitoneally with D-Luciferin (150 mg/kg), then imaged every three minutes with the auto exposure setting in supine position for a total time of 15 min. The time at which luminescence reached its maximum was used for total photon flux values (photons/sec).

For endpoint ex vivo tissue BLI quantification, mice were intraperitoneally injected with D-Luciferin (150 mg/kg), anesthetized with inhaled isoflurane, and whole body BLI was recorded 10 minutes post-luciferin injection. Following immediate euthanasia by cervical dislocation, tissues were rapidly dissected and imaged with the auto exposure setting. BLI analysis was conducted using Living Image software (v.4.7.2, PerkinElmer).

Gene expression, CRISPR-gene dependency, and metastatic potential values used for correlation analyses were sourced from the Broad Institute Dependency Map (DepMap) portal website, 24Q2 release (available at https://depmap.org/portal).

### Petal plot generation

To account for baseline activity and inter-tissue variability in our luciferase assay data, we normalized the readings against a Non-Template Control (NTC). This normalization involved dividing the raw data for each gene by the average NTC values corresponding to each tissue type. We utilized bootstrap methods for statistical reliability, resampling the normalized data to estimate mean value distributions and calculate 95% confidence intervals, thereby capturing the inherent variability in the biological data. Values indicating fold-changes above 1, suggesting normal or enhanced cell growth, were capped at 1 to focus on growth dependencies. Data visualization, inspired by Jin et al. 2020^12^, was achieved through a petal plot (a radial variant of a bar plot), succinctly depicting luciferase activities along with 95% confidence intervals. Replicates within the dataset were averaged to provide a single representative measure per tissue type. The analysis was conducted using R studio, utilizing the dplyr, ggplot2 packages for data processing and visualization, and the boot package for bootstrap confidence interval calculations.

### ^13^C-glucose tracing experiments

Infusion of [U-^13^C]-glucose (Cambridge Isotope Laboratories, CLM-1396-1) into tumor-bearing mice was performed as previously described^13,64,65^. 1E5 MDA-MB-231-Gluc cells were intracranially injected into the brain or injected into the MFP of mice and tumors were permitted to grow for 14 or 17 days, respectively. Catheters were surgically implanted into the jugular vein of tumor-bearing animals three days before the experiment and mice were fasted for the last 4 hr. [U-^13^C]-glucose was infused at a constant rate of 0.4 mg/min for 10 hr into conscious, free-moving animals, after which the animals were terminally anesthetized with infusion of Fatal Plus. Blood was collected immediately by cardiac puncture, and tumors and noncancerous tissue were collected within five min of sacrifice; for brain tumors, GFP+ tumor tissue was isolated from non-GFP+ tissue under a fluorescent dissecting microscope. Tissues were flash frozen using the BioSqueezer (BioSpec Products, 1210) and stored at −80 °C for subsequent metabolite extraction and analysis. Blood was placed into EDTA tubes (Sarstedt 41.1395.105) and centrifuged at 800 x g at 4 °C to separate plasma, which was flash frozen and stored at −80 °C. All isotope labeling experiments in mice were performed at the same time of day.

Snap frozen tissues were ground into powder using a mortar and pestle, then weighed into glass vials (Thermo Fisher Scientific, C4010-1 and C4010-60BLK). Metabolites were extracted in 1.5 mL of dichloromethane:methanol (containing 25 mg/ L of butylated hydroxytoluene (Millipore Sigma, B1378)):0.88% KCl (w/v) (8:4:3 v/v/v), vortexed for 15 min at 4 °C and centrifuged at maximum speed for 10 min at 4 °C. Polar metabolites (aqueous fraction) were transferred to Eppendorf tubes, dried under nitrogen gas and stored at −80 °C until further analysis. For plasma, 5 µL of fluid was extracted in 45 µL of 80% methanol containing 500 nM ^13^C and ^15^N amino acid standards (Cambridge Isotope Laboratory, MSK-A2-1.2). Samples were vortexed for 15 min, centrifuged at maximum speed for 10 min, and supernatant transferred to vials and analyzed by LC/MS (described below). Following analysis by LC/MS, metabolite peak areas were called using XCalibur v.2.2 (Thermo Fisher Scientific) or Compound Discoverer v.3.3 (Thermo Fisher Scientific) software. Ion counts for each metabolite were normalized to the weight of the tissue sample. Isotope corrections were applied using IsoCorrector v.3.18^66^.

### Mass spectrometry metabolite measurements

#### Quantification of metabolite levels in biological fluids

Metabolite quantification in human fluid samples was performed as described previously^4^. In brief, 5 μL of sample or external chemical standard pool (ranging from ∼5 mM to ∼1 μM) was mixed with 45 μL of acetonitrile:methanol:formic acid (75:25:0.1) extraction mix including isotopically labeled internal standards. All solvents used in the extraction mix were HPLC grade. Samples were vortexed for 15 min at 4 °C and insoluble material was sedimented by centrifugation at 16,000 x g for 10 min at 4 °C. 20 µL of the soluble polar metabolite extract was taken for LC/MS analysis (described below). Following analysis by LC/MS, metabolite identification was performed with XCalibur 2.2 software (Thermo Fisher Scientific) using a 5 ppm mass accuracy and a 0.5 min retention time window. For metabolite identification, external standard pools were used for assignment of metabolites to peaks at given m/z and retention time (see Supplementary Table 1 for the m/z and retention time for each metabolite analyzed). Absolute metabolite concentrations were determined as published^54^.

#### LC/MS analysis

Metabolite profiling was conducted on a QExactive benchtop orbitrap mass spectrometer equipped with an Ion Max source and a HESI II probe, which was coupled to a Dionex UltiMate 3000 HPLC system (Thermo Fisher Scientific). External mass calibration was performed using the standard calibration mixture every 7 days. An additional custom mass calibration was performed weekly alongside standard mass calibrations to calibrate the lower end of the spectrum (m/z 70-1050 positive mode and m/z 60-900 negative mode) using the standard calibration mixtures spiked with glycine (positive mode) and aspartate (negative mode). 2 μL of each sample was injected onto a SeQuant® ZIC®-pHILIC 150 x 2.1 mm analytical column equipped with a 2.1 x 20 mm guard column (both 5 mm particle size; EMD Millipore). Buffer A was 20 mM ammonium carbonate, 0.1% ammonium hydroxide; Buffer B was acetonitrile. The column oven and autosampler tray were held at 25 °C and 4 °C, respectively. The chromatographic gradient was run at a flow rate of 0.150 mL min^-1^ as follows: 0-20 min: linear gradient from 80-20% B; 20-20.5 min: linear gradient form 20-80% B; 20.5-28 min: hold at 80% B. The mass spectrometer was operated in full-scan, polarity-switching mode, with the spray voltage set to 3.0 kV, the heated capillary held at 275 °C, and the HESI probe held at 350 °C. The sheath gas flow was set to 40 units, the auxiliary gas flow was set to 15 units, and the sweep gas flow was set to 1 unit. MS data acquisition was performed in a range of m/z = 70–1000, with the resolution set at 70,000, the AGC target at 1×10^6^, and the maximum injection time at 20 msec.

### Cell proliferation

For assessment of proliferation by continuous live cell imaging, cells were trypsinized, pelleted and washed with PBS, counted, resuspended in the appropriate medium, then plated directly into clear 96-well plates. Cells were then treated with or without the relevant rescue metabolite, and plates were placed into an IncuCyte Live Cell Analysis Imaging System S3 (Sartorius) in a humidified incubator at 37 °C and 5% CO_2_. Images were acquired every 3 hr using the 10x objective. Cell confluence was determined from a mask generated by the IncuCyte Zoom Analysis S3 v2018B software’s standard settings. For ASNS and PYCR knockout (KO) experiments, cells were plated in DMEM. For ASS1 and PHGDH KO experiments, cells were plated in RPMI medium lacking arginine or serine, respectively. For DHODH and GART KO experiments, cells were plated in RPMI. Metabolite concentrations used for rescues: 1.15 mM arginine; 379 µM asparagine; 1 mM citrulline; 100 µM hypoxanthine; 174 µM proline; 286 µM serine; 100 µM uridine.

### Western blot

Cells were washed once in PBS and scraped into RIPA lysis buffer containing protease and phosphatase inhibitors (Cell Signaling Technology, 5871), rocked for 15 min at 4 °C, and insoluble material was sedimented by centrifugation at 21,000 x g for 10 min at 4 °C. Protein concentration was determined via Bradford Assay, and samples were mixed with LDS sample buffer (Thermo Fisher Scientific, NP0008) and 2.5% 2-mercaptoethanol then incubated at 95 °C for 5 min. Proteins were resolved by SDS-PAGE then transferred onto nitrocellulose membranes using a wet tank transfer system (Bio-Rad). Membranes were blocked in 5% milk in TBST and probed overnight at 4 °C with the appropriate antibody diluted in 5% BSA in TBST. For detection, membranes were incubated with HRP-linked anti-rabbit (Cell Signaling Technology, 7074, 1:5,000) or anti-mouse (Cell Signaling Technology, 7076, 1:5,000) IgG secondary antibody diluted in 5% milk in TBST, and chemiluminescent signal was detected using a digital imager (GE Healthcare, LAS 4000). Primary antibodies were used as follows: ASNS (Proteintech, 14681-1-AP, 1:1,000), ASS1 (Cell Signaling Technology, 70720, clone D4O4B, 1:1000), PHGDH (Sigma-Aldrich, HPA021241, 1:1000), PYCR1 (Proteintech, 13108-1-AP, 1:1000), PYCR2 (Proteintech, 17146-1-AP, 1:1000), PYCRL (Novus Biologicals, NBP2-03337, clone OTI1B12, 1:1000), PYCRL (ABclonal, A17763, 1:1000), DHODH (Santa Cruz Biotechnology, sc-166348, clone E-8, 1:100), GART (Proteintech, 13659-1-AP, 1:1000), β-actin (Cell Signaling Technology, 8457, clone D6A8, 1:5,000), and vinculin (Cell Signaling Technology, 13901, clone E1E9V, 1:1,000). In cases where re-probing of the same membrane was required, membrane was incubated with hydrogen peroxide solution 30% (w/w in water) (Sigma-Aldrich, H1009) for 1 hr at 37 °C to inactivate HRP. Western blots were quantified by densitometry using Fiji^67^; data are normalized to the loading control from the same gel.

### Lentivirus generation

LentiX cells were transfected with lentiviral packaging plasmids pMD2.G (Addgene, 12259) pMDLg/pRRE (Addgene, 12251) and pRSV-Rev (Addgene, 12253) using TransIT®-293 Transfection Reagent (Mirus, MIR 2700). 48 hr post-transfection, medium was collected and filtered through a 0.45 µm low-protein binding membrane. Lentivirus containing supernatants were used immediately or stored at −80 °C. Cell lines were transduced with virus and 10 µg/mL polybrene for 24 hr.

### CRISPR/Cas9-mediated gene knockout

Cas9-expressing cell lines were generated by infecting parental cell lines with viral particles containing lentiCas9/blast (Addgene, 52962) followed by selection with 10 µg/mL blasticidin. Two gRNAs per gene of interest were designed by CRISPick (https://portals.broadinstitute.org/gppx/crispick/public) and cloned into lentiGuide-Puro (Addgene, 52963). Primer sequences used in this study are listed in Supplementary Table 6. Cas9+ cell lines were transduced with lentiGuide-Puro-gRNA viral particles, then selected with 1 µg/mL puromycin. Single cells were plated into individual wells of 96-well plates in RPMI medium; following expansion, western blotting confirmed complete knockout. In order to generate and maintain *DHODH* or *GART* KO cells, 100 µM uridine or 100 µM hypoxanthine were supplemented in RPMI medium, respectively. For the generation of *PYCR1*, *PYCR2*, *PYCR3* (*PYCRL*) triple knockout MDA-MB-231 or HCC1806 cells, two gRNAs targeting both *PYCR1* (exons 4 and 5) and *PYCR2* (exons 2 and 3), and two gRNAs targeting *PYCR3* (exon 3), were utilized.

To knock out *Pycr1*, *Pycr2*, and *Pycr3* (*PycrI*) in EO771, we utilized ribonucleoprotein (RNP) complexes with Cas9-GFP and Cas9-RFP. The Cas9 protein and gRNAs specific to each target gene were ordered from Integrated DNA Technologies (IDT) as RNA oligonucleotides and used to assemble RNP complexes. EO771 cells were nucleofected with the RNP complexes using a 4D-Nucleofector (Lonza Bioscience) and the SF Cell Line 4D-Nucleofector X Kit L (Lonza Walkersville, V4XC2024) with Pulse Code CM-150. Post-nucleofection, cells were cultured and monitored for GFP and RFP expression. Single-cell sorting was performed 24 hr after nucleofection to isolate successfully nucleofected cells.

### Statistics and reproducibility

Sample sizes, reproducibility and statistical tests used for each figure are denoted in the figure legends. All graphs were generated using GraphPad Prism 10 (GraphPad Software).

After determining the concentration of each metabolite in each plasma or TIF sample, all multivariate statistical analysis on the data was performed using Metaboanalyst 6.0^68^. Metabolite concentrations were log-transformed and metabolites that contained greater than 50% missing values were removed prior to analysis. For the remainder of the metabolites, missing values were replaced using 1/5th of the lowest positive value. Hierarchical clustering was performed with Euclidean distance measurement and clustering by the Ward algorithm. Univariate analysis was performed comparing metabolite levels between groups where metabolite differences of interest were defined by a fold change greater than 2 and significance as a raw p-value less than 0.05 assuming unequal group variance. Using the fold change and adjusted p-value cutoffs, the number of differentially expressed metabolites were determined.

### Illustrations

Experimental schema and illustrative models were generated using BioRender (https://biorender.com/).

## Data availability

Datasets can be found in Supplementary Tables 1-6. Unprocessed western blots can be found in Supplementary Figure 1. Any additional information required to reanalyze the data reported in this paper is available from the lead contact upon reasonable request.

