## Extended Data Figure Legends for "Site of breast cancer metastasis is independent of single nutrient levels"

**Extended Data Fig. 1: Metabolite concentrations in mouse tissue interstitial fluid, plasma, and CSF.**

**a-j**, Heatmaps depicting metabolite concentrations in plasma, cerebrospinal fluid (CSF), and the indicated tissue interstitial fluids (IF) from female NOD-SCID-gamma (NSG) mice as determined by liquid chromatography/mass spectrometry (LC/MS) measurements. Metabolites are grouped according to the indicated metabolic pathways, with scale bars provided for each group to indicate the concentration ranges. Data represent the mean of n = 6 (plasma, kidney IF, liver IF, lung IF, mammary fat pad (MFP) IF, pancreas IF) or n = 4 (CSF) biological replicates. Metabolite concentrations that were below the limit of detection by LC/MS are shown in grey and labeled "nd" (not determined). **k**, Scatter plot of LC/MS measurements of metabolite concentrations (µM) in plasma or tissue IF from female NSG mice from two independent experiments. The values represent the mean of n = 4 (experiment 1) or n = 6 (experiment 2) biological replicates. The number of metabolites measured is indicated in each panel. r and p-values were determined by Pearson correlation.

**Extended Data Fig. 2: Log2 fold changes in metabolite concentrations measured between tissue fluid samples and plasma. a-b**, log2 fold change in the indicated metabolite concentrations in each tissue IF or CSF relative to plasma in female NSG mice. Data are presented as mean ± SEM and represent n = 6 (kidney IF, liver IF, lung IF, mammary fat pad (MFP) IF, pancreas IF) or n = 4 (CSF) biological replicates. **c-d**, Plots depicting the log2 fold change in polar metabolite concentrations between paired CSF and plasma samples from C57BL/6J (n = 7) or NSG (n = 4) mice, as determined by LC/MS. Data are mean ± SEM, with statistical significance assessed via one sample t-test (*p < 0.05, **p < 0.01, ***p < 0.001, ****p < 0.0001).

**Extended Data Fig. 3: Assessment of amino acid and nucleotide synthesis pathways in breast cancer cell lines.**

**a**, Schematic illustrating the amino acid or nucleotide de novo synthesis pathways highlighting the enzymes targeted in this study to generate various auxotrophs. Genes encoding the enzymes knocked out in breast cancer cells are highlighted in red, and relevant metabolites that are synthesized are highlighted in blue. Asp: aspartate; DHO: dihydroorotate; Glu: glutamate; Gln: glutamine; IMP: inosine monophosphate; P5C: 1-pyrroline-5-carboxylic acid; PRPP: phosphoribosyl diphosphate; UMP: uridine monophosphate. **b**, Western blot analysis of parental MDA-MB-231, EO771, and HCC1806 breast cancer cells for expression of the indicated proteins. **c**, Quantification of the western blots in (b). The signal intensity of each indicated protein is normalized relative to the vinculin loading control. Data are presented as mean ± SD (n = 3 biological replicates). Statistical analysis was performed using an ordinary one-way ANOVA with Holm-Sidak's multiple comparisons test (*p < 0.05, **p < 0.01, ****p < 0.0001).

**Extended Data Fig. 4: Validation of auxotroph cell lines by western blot.**

**a-f**, Western blot analysis of non-targeting control (Ctrl) or knockout (KO) auxotroph clonal cell lines for expression of the indicated proteins in MDA-MB-231, HCC1806, or EO771 cells. PYCR KO represents PYCR1, 2, and 3 triple KO.

**Extended Data Fig. 5: Validation of auxotroph cell lines by assessing proliferation with or without rescue metabolites.**

**a-f**, Percent confluence of the indicated control (Ctrl) or knockout (KO) clonal cell lines over time when cultured in medium with or without the relevant rescue metabolites. For ASNS KO (a), cells were cultured in DMEM. For ASS1 KO (b) and PHGDH KO (c) experiments, cells were cultured in RPMI medium lacking arginine or serine, respectively. For PYCR KO experiment (d), cells were cultured in DMEM. For DHODH KO (e) and GART KO (f) experiments, cells were cultured in RPMI. Metabolite concentrations used for rescues: 1.15 mM arginine; 379 µM asparagine; 1 mM citrulline; 100 µM hypoxanthine; 174 µM proline; 286 µM serine; 100 µM uridine. Data are mean ± SD and represent n = 3 biological replicates from a representative experiment performed independently at least twice. Asn: asparagine; Arg: arginine; Cit: citrulline; Ser: serine; Pro: proline; Urd: uridine; Hx: hypoxanthine. PYCR KO represents PYCR1, 2, and 3 triple KO.

**Extended Data Fig. 6: Assessment of breast cancer cell metastasis in mice.**

**a**, Whole body bioluminescence imaging (BLI) in units of total flux (photons/sec) of MDA-MB-231-Fluc control, HCC1806-Fluc control, and EO771-Fluc control cell lines over time following intracardiac injection into female NSG or female C57BL/6J mice. Data are mean ± SEM and represent n = 7 NSG mice (MDA-MB-231), n = 5 NSG mice (HCC1806), or n = 7 C57BL/6J mice (EO771). Data are from the same experiments presented in Extended Data Fig. 7c (MDA-MB-231), 7i (HCC1806), and 8c (EO771). **b**, BLI and quantification of metastasis burden in the organs of mice receiving intracardiac injection of indicated cell lines as described in (a). **c**, Representative images of mice and tissues displaying tumor burden resulting from intracardiac injection as described in (a). Color scales reflect BLI radiance and min and max are noted to the right of each set of images.

**Extended Data Fig. 7: Metastasis of MDA-MB-231 and HCC1806 auxotrophs following intracardiac injections in mice.**

**a-l**, Left panels, whole body bioluminescence imaging (BLI) in units of total flux (photons/sec) tracking the progression of metastasis over time in female NSG mice with either MDA-MB-231-Fluc (a-f) or HCC1806-Fluc (g-l) auxotroph or control (ctrl) cells as indicated. The final whole body BLI points represent the endpoint when all tissues were harvested. Right panels, quantitative analysis of tissue-specific BLI, illustrating the relative change in metastatic burden across various organs following intracardiac injections of ctrl versus auxotroph cells. Depletion values represent the average fold change in metastatic load in the organs injected with auxotroph cells relative to ctrl cells (◦: <2-fold depletion; +: 2-to 5-fold depletion; ++: 5 to 10-fold depletion; +++: >10-fold depletion). Data are presented as mean ± SEM and the number of mice per experimental group is indicated within each panel; two bones, kidneys, or ovaries were analyzed per mouse. Statistical analysis was performed using an unpaired t-test with Holm-Sidak multiple comparisons and Welch correction for significance (*p < 0.05, **p < 0.01, ***p < 0.001, ****p < 0.0001). PYCR KO represents PYCR1, 2, and 3 triple KO.

**Extended Data Fig. 8: Metastasis of EO771 auxotrophs following intracardiac injections in mice.**

**a-f**, Left panels, whole body bioluminescence imaging (BLI) in units of total flux (photons/sec) tracking the progression of metastasis over time in female C57BL/6J mice with EO771-Fluc auxotroph or control (ctrl) cells as indicated. The final whole body BLI points represent the endpoint when all tissues were harvested. Right panels, quantitative analysis of tissue-specific (BLI), illustrating the relative change in metastatic burden across various organs following intracardiac injections of ctrl versus auxotroph cells. Depletion values represent the average fold change in metastatic load in the organs injected with auxotroph cells relative to ctrl cells (◦: <2-fold depletion; +: 2-to 5-fold depletion; ++: 5 to 10-fold depletion; +++: >10-fold depletion). Data are presented as mean ± SEM and the number of mice per experimental group is indicated within each panel; two bones, kidneys, or ovaries were analyzed per mouse. Statistical analysis was performed using an unpaired t-test with Holm-Sidak multiple comparisons and Welch correction for significance (**p < 0.01, ****p < 0.0001). PYCR KO represents PYCR1, 2, and 3 triple KO.

**Extended Data Fig. 9: Correlations of auxotroph metastatic potential with tissue metabolite levels.**

**a-b**, Scatter plots correlating the concentration of citrulline (a) or trans-uroconate (b) in tissue interstitial fluids with the dependency of MDA-MB-231 cells on ASNS (a) or ASS1 (b) for metastatic growth in that tissue. The x-axis shows the tissue metabolite concentration, while the y-axis displays the dependency as a log2 fold change of knockout (KO) compared to control (Ctrl). Symbols denote the metabolite concentration in specific tissues, with brain values derived from CSF measurements. Data represent mean ± SEM, and the raw data used to derive dependency values involve the number of mice per experimental group presented in Extended Data Fig. 7-8. Pearson correlation coefficients (r) and p-values are provided to assess statistical significance. **c**, Volcano plots depicting the Pearson correlation values and p-values for the relationships between metabolite levels and metastatic potential of auxotroph cell lines following intracardiac injection. Pearson correlation values and p-values are derived as in the representative scatter plots in (a-b). **d**, Bar plots representing the fraction of metabolites that exhibit significant positive (pos) or negative (neg) correlations with the metastatic potential of the indicated auxotroph cell lines from (c).

**Extended Data Fig. 10: Tumor growth of auxotroph versus control cells following intracranial or MFP implantations.**

**a**, Tumor growth in NSG mice injected in the brain (upper panels) and mammary fat pad (MFP) (lower panels) with control (Ctrl) or knockout (KO) MDA-MB-231-Gluc (a-f) or HCC1806-Gluc (g-l) cell lines. Tumors were monitored over time using secreted Gaussia luciferase (Gluc) as a marker for tumor abundance. Data are presented as mean ± SEM and the number of mice per experimental group is indicated within each panel. Statistical analysis was performed using a two-way ANOVA across timepoints and groups and significance values are indicated within each panel. **m-r**, Tumor growth in C57BL/6J mice injected in the brain with Ctrl or KO EO771-Gluc cell lines. Tumors were monitored over time using secreted Gaussia luciferase (Gluc) as a marker for tumor abundance. Data are presented as mean ± SEM and the number of mice per experimental group is indicated within each panel. Statistical analysis was performed using a two-way ANOVA across timepoints and groups and significance values are indicated within each panel. PYCR KO represents PYCR1, 2, and 3 triple KO.

**Extended Data Fig. 11: Mouse survival following intracranial implantations of auxotroph versus control cells.**

**a-f**, Upper and middle panels, Kaplan-Meier survival curves for NSG mice bearing MDA-MB-231-Gluc or HCC1806-Gluc control (ctrl) or knockout (KO) brain tumors. Lower panels, Kaplan-Meier survival curves for C57BL/6J mice bearing ctrl or KO EO771-Gluc brain tumors. Data are from the experiments presented in Extended Data Fig. 10. Number of mice in each group are noted in each figure panel. Statistical analysis was performed using Kaplan-Meier log-rank (Mantel-Cox) test and significance values are indicated within each panel. **g**, Analysis of the relative dependency of metabolite auxotroph cell lines on specific metabolic genes to grow in the brain, expressed as log2 fold change of KO relative to ctrl following intracranial or intracardiac injection methods. Data are mean ± SEM. The number of mice per experimental group are indicated in Extended Data Fig. 7-8 and Extended Data Fig. 10. PYCR represents PYCR1, 2, and 3 triple KO.

**Extended Data Fig. 12: Metabolite levels in MDA-MB-231 brain or MFP tumors.**

**a**, Tumor growth in NSG mice injected in the mammary fat pad (MFP) or brain or with MDA-MB-231-Gluc cells. Tumors were monitored over time using secreted Gaussia luciferase (Gluc) as a marker for tumor abundance. Data points represent mean ± SEM for n = 5 (MFP tumor) or n = 4 (brain tumor) biological replicates. At the experimental endpoint, following [U-^13^C]-glucose infusion, both tumors and their corresponding noncancerous control tissues were harvested from the same mice (MFP tumor with adjacent MFP tissue, and brain tumor with adjacent brain tissue). **b**, Normalized peak area values for indicated metabolites measured in MFP, MFP tumor, brain, or brain tumor tissue isolated from the mice shown in (a) at endpoint. **c**, Isotopolog distribution showing fractional labeling of the indicated metabolites as measured by LC/MS in cancerous tissues (MDA-MB-231 tumors in the brain and MFP) and corresponding noncancerous tissues (brain and MFP) from NSG mice infused with [U-^13^C]-glucose. **d**, Normalized peak area values for the indicated metabolites measured in MFP, MFP tumor, brain, or brain tumor isolated from the mice shown in (a) at endpoint. **e**, Isotopolog distribution showing fractional labeling of the indicated metabolites as measured by LC/MS in cancerous tissues (MDA-MB-231 tumors in the brain and MFP) and corresponding noncancerous tissues (brain and MFP) from NSG mice infused with [U-^13^C]-glucose. In b-e, data points represent mean ± SEM for n = 5 (MFP tumor, MFP), 3 (noncancerous brain), and 4 (brain tumor) biological replicates.

**Extended Data Fig. 13: Assessment of nucleotides in MDA-MB-231 brain or MFP tumors.**

**a**, Normalized peak area values measured for the indicated metabolites in MFP, MFP tumor, brain, or brain tumor tissue isolated from female NSG mice. **b**, Isotopolog distribution showing fractional labeling of the indicated metabolites as measured by LC/MS in cancerous tissues (MDA-MB-231 tumors in the brain and MFP) and corresponding noncancerous tissues (brain and MFP) from NSG mice infused with [U-^13^C]-glucose. In a-b, data points represent mean ± SEM for n = 5 (MFP tumor, MFP), 3 (noncancerous brain), and 4 (brain tumor) biological replicates.

**Extended Data Fig. 14: Correlating gene expression with tissue-specific metastatic potential.**

**a-e**, Scatter plots correlating the metastatic potential of breast cancer cells to the brain (a), liver (b), lung (c), kidney (d), and bone (e) with RNA expression (transcript per million, TPM) of the indicated genes reported in the Dependency Map portal. Each dot represents a cell line and p-values derived from Pearson correlations are indicated within each plot.

**Extended Data Fig. 15:** **Correlating gene dependency with tissue-specific metastatic potential.**

**a-d**, Scatter plots correlating the metastatic potential of breast cancer cell lines to the brain (a), liver (b), kidney (c), and bone (d) with in vitro CRISPR dependency of the indicated genes as reported in the Dependency Map portal. Each dot represents a cell line and p-values derived from Pearson correlations are indicated within each plot.
